# Stochastic Growth Modeling of Vascular Plaque Dynamics and Derivation of Optimal Dosing Curves

**DOI:** 10.64898/2026.05.30.728429

**Authors:** Taiga Kadowaki, Atsushi Tero

## Abstract

Targeted drug delivery offers a promising approach for personalized medicine in treating vascular stenosis. However, biomechanical constraints, such as drug washout by high-velocity central blood flow and unintended absorption by healthy vascular walls, complicate the determination of optimal dosing locations. Conventional three-dimensional computational fluid dynamics (CFD) provides precise flow analysis but incurs prohibitive computational costs, making long-term tracking of plaque growth and reverse-engineering of optimal delivery highly inefficient. In this study, we propose a pseudo-3D stochastic growth model that dramatically reduces computational load while capturing the essential dynamics of plaque progression and regression. By modeling the advection-diffusion of lipid and drug particles as a discrete Markov process within a Stokes flow field, we simulate the morphological evolution of plaques under continuous and interrupted targeted therapies. Furthermore, by formulating the drug transport process as an absorbing Markov chain with boundaries at the healthy walls and vessel outlet, we calculate the exact reaching probability and mean first passage time (MFPT) to the plaque. Based on these probability distributions, we discover continuous “Optimal Dosing Curves”, which indicate the most effective spatial coordinates for catheter-based drug release to maximize therapeutic efficacy. This mathematical framework not only elucidates the stochastic nature of vascular plaque dynamics but also provides a scalable, computationally efficient foundation for optimizing targeted drug delivery in personalized medicine.

## 1. Introduction

Vascular stenosis, driven by progressive arterial plaque accumulation, is a leading cause of life-threatening cardiovascular diseases such as myocardial infarction and ischemic stroke. Targeted drug delivery via catheter-based micro-injection offers a promising approach to enhance efficacy while minimizing systemic toxicity [1]. However, its clinical optimization is highly constrained by biomechanical competition: therapeutic particles are often washed out by high-velocity blood flow or prematurely absorbed by healthy endothelial walls. Determining the optimal injection coordinates to overcome these transport barriers is therefore essential for personalized medical intervention.

Traditional three-dimensional computational fluid dynamics (CFD) provides precise hemodynamic insights but incurs prohibitive computational costs. This intensive overhead makes standard fluid models ill-suited for tracking long-term plaque morphological evolution or performing the iterative inverse-engineering required to locate optimal drug delivery positions. Consequently, there is an urgent need for an efficient mathematical framework that captures essential physical transport mechanics without heavy computational burdens.

To bridge this methodological gap, this study introduces a pseudo-three-dimensional stochastic growth model integrated with an absorbing Markov chain. Crucially, while real pathological networks possess complex, multi-branched structures, establishing a robust mathematical foundation for a single vascular segment (edge) is an indispensable first step before scaling up to full networks. By applying the lubrication approximation to the Stokes flow field and mapping advection-diffusion transport onto a discrete spatial grid, we drastically reduce computational load while strictly preserving mass conservation laws. This probabilistic framework enables the synchronous simulation of plaque progression and regression, allowing for the direct computation of reaching probabilities and mean first passage times (MFPT) within a single edge segment.

The remainder of this paper is organized as follows: Section 2 formulates the mathematical framework and model assumptions; Section 3 presents numerical results demonstrating plaque morphological evolution and the derivation of optimal dosing curves; Section 4 discusses biomedical relevance and clinical implications; and Section 5 concludes the paper with directions for future network-scale research.

## 2. Mathematical Modeling

### 2.1. Conceptual Framework and Model Assumptions

To bridge micro-scale particle transport and long-term plaque evolution, we model an isolated segment of a coronary or cerebral artery as a single edge within a larger vascular network. For mathematical generality, the flow and transport domain is formulated within a dimensionless, axially symmetric spatial coordinate system.

The healthy vessel is treated as a straight, rigid cylindrical tube defined by its total longitudinal length *L* and uniform initial radius *R*. Under axial symmetry, the system reduces to a 2D computational grid in (*z, r*), where *z* ∈ [0, *L*] is the axial position and *r* ∈ [0, *R*] is the radial distance. A localized plaque of length *L*_*p*_ is centered at *z*_*c*_, restricting the flow space with a dynamically changing effective radius *r*_*p*_(*z, t*).

**Figure 1:**
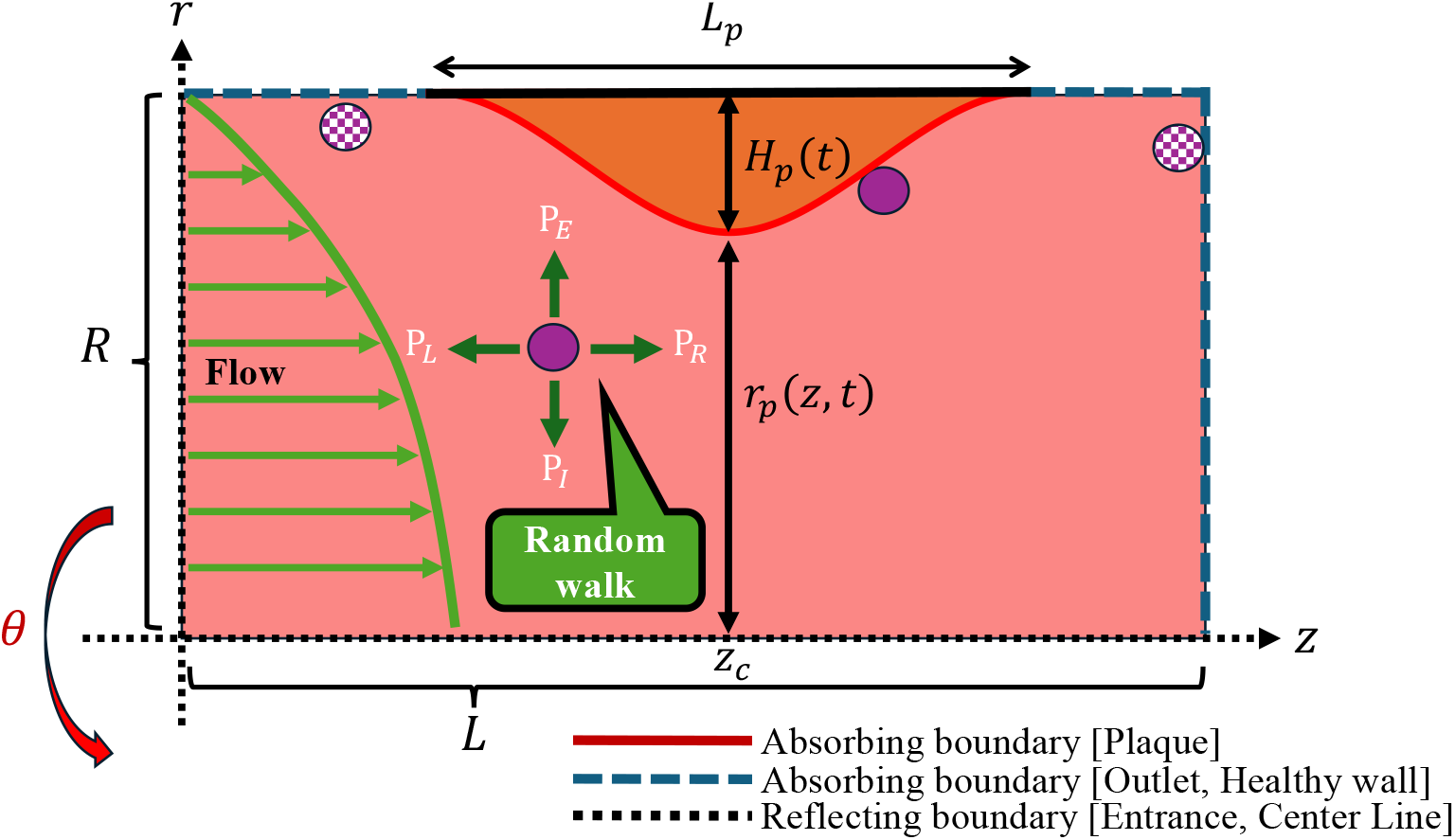
Schematic illustration of the proposed mathematical framework. The three-dimensional vascular segment of length *L* and radius *R* is simplified into an axially symmetric two-dimensional computational domain. The model couples (i) micro-scale stochastic transport of lipid and drug particles, (ii) mid-scale fluid dynamics via the lubrication approximation, and (iii) macro-scale morphological evolution of the plaque of length *L*_*p*_ centered at *z*_*c*_ based on the net volume change of absorbed particles.

The following assumptions are fundamental to our model:

#### 1. Idealized Geometry

The vascular segment is treated as a straight cylindrical tube defined by length *L* and radius *R*. All morphological changes are assumed to be axially symmetric, allowing the system to be reduced to a 2D (*z, r*) grid for computational efficiency.

#### 2. Quasi-Steady Flow

While the plaque updates its height at a macroscale time interval, the blood flow is assumed to be in a steady laminar state at each update step, neglecting pulsatile cardiac cycles.

#### 3. Attachment-Based Growth

Plaque progression and regression are driven solely by the cumulative number of attached particles. A single-particle attachment event is mapped to a localized volume change that maintains the idealized cosine-shaped profile of the plaque defined within the region *L*_*p*_ around the center *z*_*c*_.

#### 4. Absorbing Boundaries

Throughout all evolution and dosing stages, healthy endothelial walls and the vessel outlet (*z* = *L*) are defined as absorbing boundaries to strictly evaluate transport efficiency against unintended particle loss.

### 2.2. Fluid Dynamics and Lubrication Approximation

Consistent with the assumptions of an idealized cylindrical geometry and a quasi-steady laminar state, the local blood flow field within the stenosed domain can be determined analytically. Under low Reynolds number conditions typical of microvascular domains, viscous forces dominate inertial forces, reducing the Navier-Stokes equations to the Stokes equations. Furthermore, because the longitudinal scale of the vessel segments is sufficiently larger than its radial scale, we apply the lubrication approximation to the fluid field [2].

Assuming a steady and incompressible laminar flow, the axial velocity *v*_*z*_(*z, r*) and radial velocity *v*_*r*_(*z, r*) at a given coordinate are explicitly derived as follows:

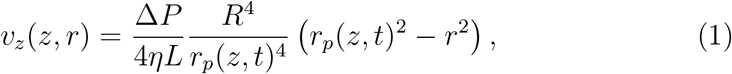

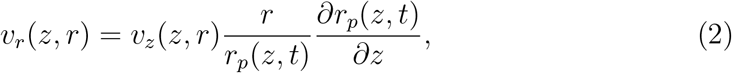

where Δ*P* is the pressure drop across the vessel segment, and *η* is the dynamic viscosity of the blood. Equation (1) represents a modified Poiseuille flow field that scales dynamically with the changing shape of the lumen boundary. The radial velocity component in Eq. (2) is dictated by the fluid continuity equation, ensuring that the total volumetric flow rate strictly satisfies the mass conservation law at every position along the axial direction.

### 2.3. Stochastic Particle Transport Model

The transport of blood lipids and drug particles is governed by the advection-diffusion equation. To handle long-term plaque evolution and reverse-engineer delivery efficiency with minimal computational cost, we transform the continuous governing equation into a discrete random walk process based on a discrete master equation [3].

By setting both the spatial grid size Δ*z* = Δ*r* = 1 and the temporal step Δ*t* = 1, the isotropic diffusion coefficient is naturally fixed at *D* = 1*/*4. The continuous advection-diffusion equation for the particle concentration *u*(*z, r, t*) is expressed as:

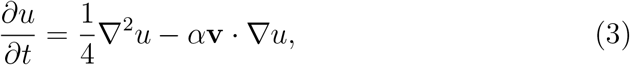

where **v** = (*v*_*z*_, *v*_*r*_) is the velocity field defined in Eqs. (1) and (2). The parameter *α* is a dimensionless scaling constant that aggregates various physical properties of the particles, including their size, shape, and buoyancy. Physically, *α* corresponds to the local Peclet number (*Pe*), adjusting the relative dominance of advective transport over diffusion while the diffusion rate remains fixed.

On the discrete two-dimensional grid, the motion of a particle at position **x** = (*z, r*) is determined by local transition probabilities derived from the discrete stochastic framework [3]. Reflecting the cylindrical coordinate system and the flow field, the transition probabilities to the adjacent grid cells—outer radial (*P*_*E*_), inner radial (*P*_*I*_), left axial (*P*_*L*_), and right axial (*P*_*R*_)—are defined as:

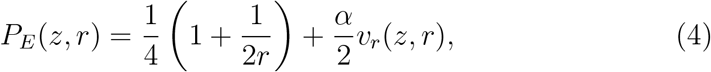

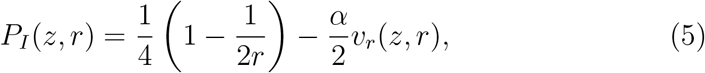

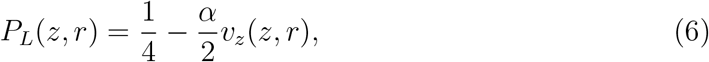

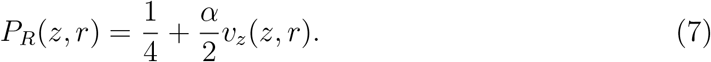

These probabilities strictly satisfy the normalization condition *P*_*E*_ + *P*_*I*_ +*P*_*L*_ + *P*_*R*_ = 1 at every interior grid point.

### 2.4. Plaque Morphological Evolution (Macro Growth Model)

While the discrete random walk governs the micro-scale transport of individual particles, the macro-scale structural evolution of the vascular plaque is modeled by assuming an idealized geometry that updates based on the cumulative number of attached particles. We assume that the plaque maintaining an axially symmetric, smooth cosine-shaped profile. When a stenosis is present within the axial range *z* ∈ [*z*_*c*_ −*L*_*p*_*/*2, *z*_*c*_ + *L*_*p*_*/*2], where *z*_*c*_ is the center of the lesion and *L*_*p*_ is the longitudinal plaque length, the effective lumen radius *r*_*p*_(*z, t*) is geometrically defined as:

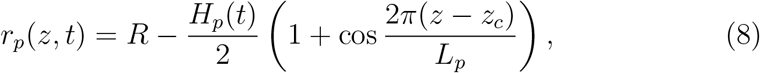

where *R* is the initial healthy vessel radius, and *H*_*p*_(*t*) represents the peak height of the plaque at time *t*. For the axial regions outside this stenotic zone, the radius remains constant (*r*_*p*_ = *R*).

The dynamic progression and regression of the plaque are driven by the volumetric contribution of attached particles. Let *N*_*f*_ (*t*) and *N*_*d*_(*t*) be the cumulative number of attached lipid (form) particles and drug (destruction) particles at time *t*, respectively. Assuming that the total volume change of the plaque is proportional to the net volume of these absorbed particles, the temporal increment of the peak height Δ*H*_*p*_(*t*) per unit time step is derived from the volumetric conservation law as follows:

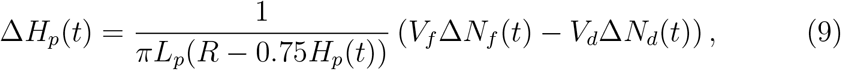

where *V*_*f*_ and *V*_*d*_ represent the discrete volume equivalents of a single lipid and drug particle, respectively.

This formulation establishes a pseudo-three-dimensional macro growth model. The peak height *H*_*p*_(*t*) is updated at each macro time step using Eq. (9), which directly modifies the effective vascular radius *r*_*p*_(*z, t*) via Eq. (8). This updated geometry is then immediately fed back into the fluid dynamics equations Eqs. (1) and (2), altering the local velocity field and the subsequent particle transition probabilities. This continuous feedback loop allows us to track the nonlinear growth and regression dynamics within a highly efficient computational framework.

### 2.5. Absorbing Markov Chain for Targeted Dosing

To evaluate the therapeutic efficiency of targeted drug delivery, we extend the stochastic framework to an open system using an absorbing Markov chain. In this clinical scenario, drug particles released into the flow domain experience competing biomechanical destinations, which can be effectively analyzed using the classical theory of first-passage processes [4].

We define the plaque boundary as the target destination. Conversely, the healthy vascular walls (*r* = *r*_*p*_(*z*) where no plaque exists) and the vessel outlet (*z* = *L*) are treated as absorbing boundaries. In this model, the adhesion efficiency of drug particles hitting the vascular wall or the plaque is assumed to be 100%, meaning that any particle touching these boundaries is immediately absorbed and removed from the active flow field.

Let *T*_(*z,r*)_(*H*_*p*_) be the Mean First Passage Time (MFPT) representing the expected number of steps for a particle injected at grid position (*z, r*) to reach any absorbing boundary, given the plaque peak height *H*_*p*_. Similarly, let Π_(*z,r*)_(*H*_*p*_) be the reaching probability that a particle injected at (*z, r*) successfully reaches the plaque rather than the healthy wall or the outlet. Since the local flow field and subsequent transition probabilities dynamically change depending on the plaque geometry, both values are fundamentally functions of *H*_*p*_. For notational simplicity, we omit the function argument and write *T*_(*z,r*)_ and Π_(*z,r*)_ when *H*_*p*_ is fixed.

The relationship between transport geometry and first-passage kinetics allows us to directly compute these values in confined flow domains [5]. By treating the grid domain as a system of linear equations, *T*_(*z,r*)_ and Π_(*z,r*)_ can be computed directly using the following grid-based difference equations:

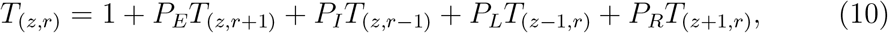

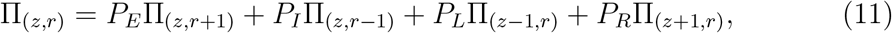

subject to the boundary conditions:

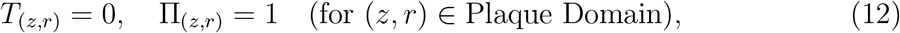

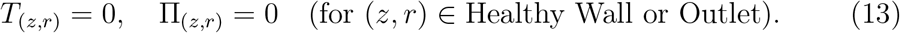

By solving this grid-based linear system over the entire flow domain, we can comprehensively map the spatial distribution of drug delivery efficiency and identify the coordinates that maximize Π_(*z,r*)_, which defines the optimal dosing curve.

## 3. Simulation Results

In this section, we present numerical simulation results based on the stochastic particle transport and plaque evolution model formulated in Section 2. We first demonstrate the temporal morphological dynamics of the plaque under competing particle attachments. Subsequently, we evaluate the spatial transport efficiency in an open vascular system to derive the optimal dosing curves for catheter-based interventions. The baseline spatial and temporal steps are fixed at Δ*z* = Δ*r* = 1 and Δ*t* = 1, respectively.

### 3.1. Plaque Growth and Regression Dynamics

We first investigate the long-term structural changes of the plaque within the vascular segment. To maintain physical consistency with the targeted dosing analysis, all healthy endothelial walls are uniformly treated as absorbing boundaries where lipid and drug particles are eliminated upon contact. This setup allows us to track the realistic morphological evolution driven by the competing accumulation of particles at the stenotic lesion, while naturally accounting for the continuous particle loss along the healthy walls.

For this temporal simulation, the dimensionless vessel geometry is configured with a total length *L* = 600 and initial radius *R* = 150. The initial plaque is centered at *z*_*c*_ = 300 with a longitudinal length *L*_*p*_ = 200 and an initial peak height *H*_*p*_(0) = 40. The fluid field is driven by a pressure gradient of 0.00005 with unit viscosity (*η* = 1.0). To simulate competitive growth under generalized systemic circulation, lipid and drug particles (with advection sensitivity *α* = 1.0) are continuously introduced into the domain exclusively from the upstream boundary (*z* = 0) at rates of 20 and 10 particles per time step, respectively. The macroscopic plaque geometry is updated every 100 time steps based on a volume equivalent *V*_*f*_ = *V*_*d*_ = 500 per attached particle. When lipid particles (form particles) dominantly enter the flow domain, they attach to the plaque surface, causing the plaque peak height *H*_*p*_(*t*) to increase over time. As the plaque grows and the effective radius *r*_*p*_(*z, t*) narrows, the local flow velocity significantly accelerates at the stenotic throat due to the mass conservation law under the lubrication approximation. Conversely, when drug particles (destruction particles) are introduced, they adhere to the plaque and induce regression, reducing *H*_*p*_(*t*) and restoring the cross-sectional area of the lumen.

**Figure 2:**
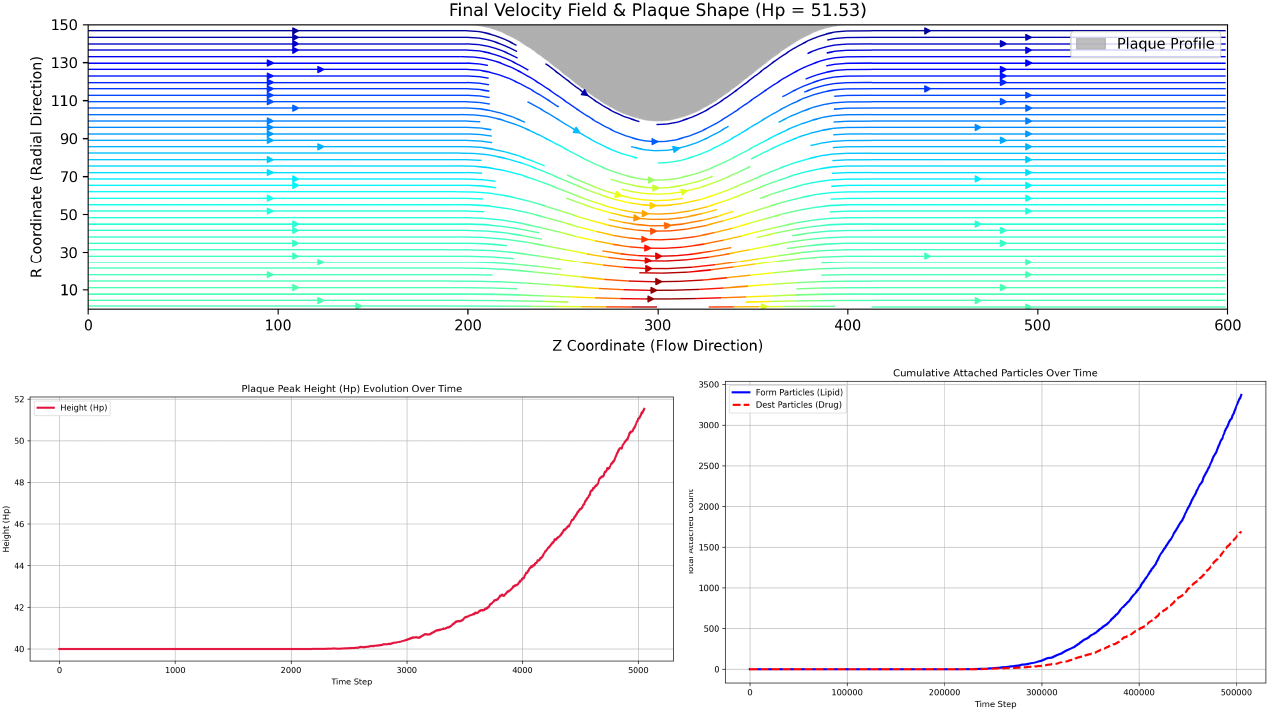
Numerical simulation of plaque growth and regression dynamics accounting for particle loss at healthy endothelial walls. The upper panel illustrates the streamlined flow field and the deformed plaque profile at the final simulation step. The lower-left plot shows the temporal evolution of the plaque peak height (*H*_*p*_), and the lower-right plot indicates the cumulative number of attached lipid (form) and drug (destruction) particles over time.

### 3.2. Targeted Dosing Analysis in Open Systems

Next, we evaluate the particle transport efficiency in an open vascular system, where healthy endothelial walls and the vessel outlet act as absorbing boundaries. The fundamental geometric and fluid parameters remain identical to the previous scenario (*L* = 600, *z*_*c*_ = 300).

By solving the linear grid-based system defined in Eqs. (10) and (11), we obtain the global spatial distributions of the Mean First Passage Time (MFPT) and the reaching probability *Q*_(*z,r*)_ across the entire flow domain.

The spatial distribution of MFPT reveals the expected time for a particle to hit any absorbing boundary from its initial injection coordinate. Particles injected near the high-velocity centerline in the upstream region exhibit relatively short MFPTs because they are quickly transported downstream by the advection-dominant flow. On the other hand, particles injected near the walls or deep inside the upstream boundary take a longer time to reach the plaque due to the slower velocity field near the boundaries, which increases the time spent in diffusion.

**Figure 3:**
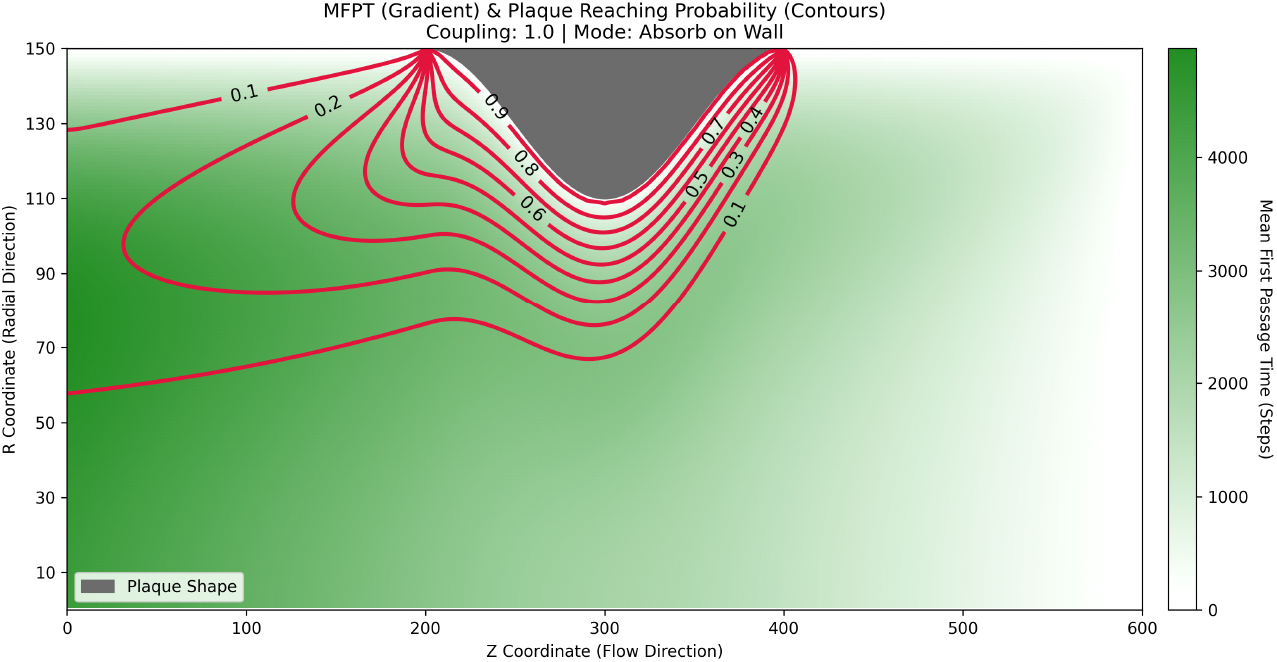
Spatial distribution of the Mean First Passage Time (MFPT) and contour map of the plaque reaching probability *Q*_(*z,r*)_ in an open vascular domain with absorbing walls. The color gradient represents the expected number of transport steps (MFPT), while the solid red lines indicate the contours of the probability of successfully reaching the plaque surface.

To investigate the optimal spatial trajectory from a broader upstream perspective, we extend the domain to *L* = 3000 and shift the plaque center to *z*_*c*_ = 2700. Here, we model targeted drug delivery via an ideal micro-catheter with negligible thickness (*r* = 0), ensuring the local Stokes flow field remains undisturbed.

By tracking the radial maximum of the reaching probability Π_(*z,r*)_(*H*_*p*_) at each axial position *z*, we extract a continuous spatial trajectory of peak targeting efficiency. We define this “Optimal Dosing Curve” as:

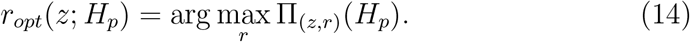

Far upstream, where plaque-induced flow disturbance is negligible, the optimal dosing curve strictly aligns with the vessel centerline (*r* = 0) to minimize premature absorption by healthy walls. However, as flow approaches the stenosis, the optimal curve dynamically bends toward the plaque surface. Furthermore, if clinical constraints restrict the catheter strictly to the centerline (*r* = 0), the optimal axial release position *z*_*opt*,0_ and its corresponding maximum reaching probability 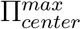 are evaluated as:

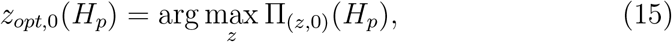

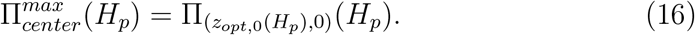

This bending behavior and centerline efficiency metric represent the optimal trade-off between advective washout and wall absorption, providing a precise spatial guideline for catheter-based interventions.

**Figure 4:**
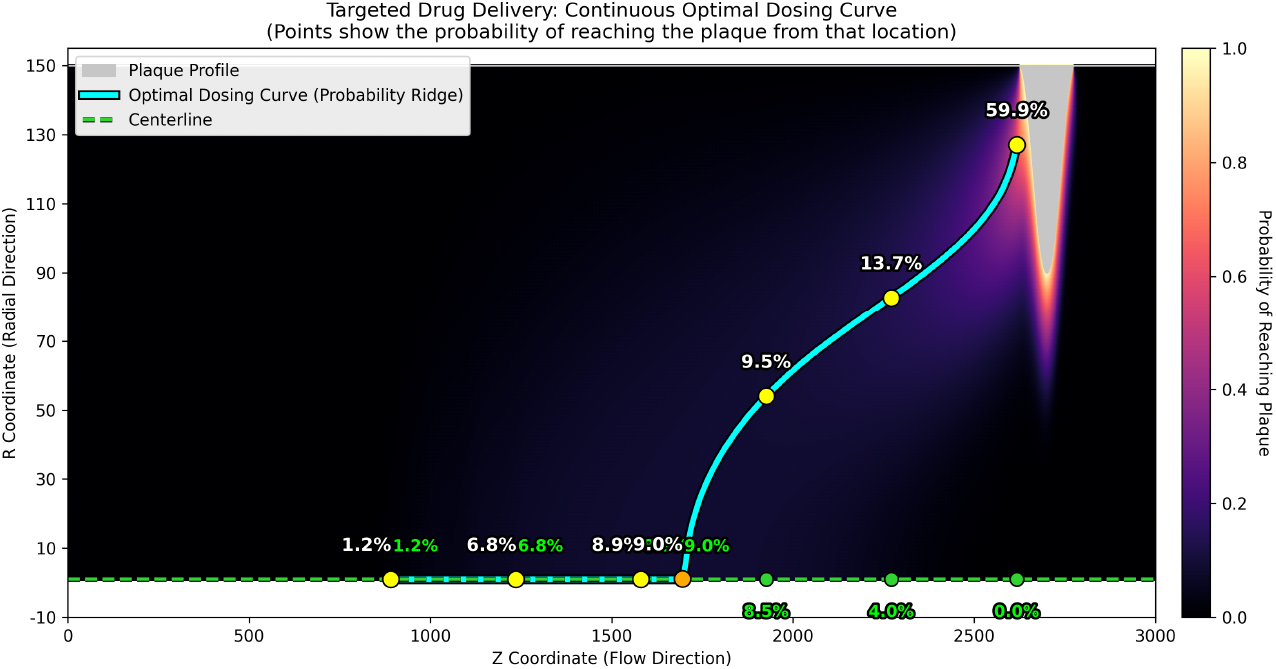
The continuous Optimal Dosing Curve derived from the probability ridge within the extended open vascular domain (*L* = 3000). The cyan solid line indicates the spatial trajectory that maximizes the plaque reaching probability from each axial position *z*. The dashed green line represents the vessel centerline, highlighting how the optimal injection path deviates from the center as it approaches the stenotic lesion to overcome the flow disturbance.

Notably, from a geometric perspective, the optimal axial position *z*_*opt*,0_(*H*_*p*_) under the centerline constraint precisely corresponds to the intersection (or bifurcation point) where the continuous optimal dosing curve *r*_*opt*_(*z*; *H*_*p*_) begins to deviate from the vessel centerline. Downstream of this intersection, the spatial ridge of maximum probability shifts decisively toward the plaque surface. This implies that releasing drug particles along the centerline further downstream of this point results in a rapid decline in targeting efficiency due to accelerated central advective washout. Therefore, this specific intersection point represents the absolute best injection coordinate available under centerline-restricted catheter operations.

## 4. Discussion

### 4.1. Bioengineering and Clinical Relevance

The primary contribution of the proposed stochastic framework is its ability to rapidly evaluate targeted drug delivery efficiency while considering complex individual vascular geometry. In the clinical management of vascular stenosis, determining the spatial entry point for catheter-based drug release has traditionally relied on the visual judgment and empirical intuition of clinicians. By calculating the exact reaching probability Π_**x**_ and mapping the continuous optimal dosing curves, our model provides an objective, geometry-driven guideline to maximize therapeutic concentration at the pathological plaque site.

Furthermore, the mathematical approach using a pseudo-three-dimensional model and an absorbing Markov chain offers an overwhelming advantage in terms of computational efficiency compared to conventional 3D CFD models. While traditional fluid-structure interaction simulations require hours or days of supercomputing resources, our grid-based linear system can be solved almost instantaneously. This low computational cost is highly valuable for real-time, bedside clinical decision-making, allowing medical practitioners to perform patient-specific inverse-engineering and optimize catheter placement during or immediately prior to surgical interventions.

From a safety perspective, minimizing drug washout into the general circulation and avoiding unintended absorption by healthy endothelial walls are crucial for reducing systemic toxicity and local side effects. Our model explicitly accounts for these competing biomechanical sinks by treating healthy walls as absorbing boundaries with 100% adhesion efficiency. The derived optimal dosing curves demonstrate that the safest injection path does not always correspond to the geometric centerline of the vessel. As the stenosis advances, the curve dynamically shifts, providing a bioengineering solution to bypass the intensified wall-shear stress and central convective flow that cause targeting failure.

### 4.2. Application to General Particle Accumulation Systems

Beyond its immediate application to vascular medicine, the stochastic interface growth and advection-diffusion framework developed in this study can be scaled to analyze general fluid-solid interaction systems. The physics of particles traveling through a confined domain and accumulating on a boundary surface is shared across various engineering fields.

For instance, this model can be effectively applied to analyze mineral precipitation and scale formation inside industrial pipe systems, such as water heaters and heat exchangers. In such systems, the continuous deposition of calcium carbonate or other minerals narrows the effective pipe radius, accelerates local flow velocity, and alters the transport probability of subsequent mineral particles. By replacing the dynamic properties of therapeutic drug particles with mineral saturation factors, our discrete master equation approach can track long-term scale growth dynamics and predict clogging locations with significantly lower computational overhead than standard multiphase fluid simulations.

## 5. Conclusion

In this paper, we developed a computationally efficient, pseudo-three-dimensional stochastic growth model combined with an absorbing Markov chain to simulate vascular plaque dynamics and optimize targeted drug delivery. By employing the lubrication approximation to simplify the Stokes flow field and transforming the continuous advection-diffusion process into local transition probabilities, we successfully captured the essential physics of particle transport without incurring heavy computational costs. Our dynamic simulations successfully reproduced the progressive growth and drug-induced regression of arterial plaques over time, even when accounting for continuous particle loss at healthy endothelial walls.

Moreover, by treating these healthy walls and the vessel outlet as absorbing boundaries, we comprehensively mapped the spatial distribution of the mean first passage time and derived continuous optimal dosing curves that represent the peak targeting efficiency. Crucially, we discovered that under the clinical constraint of centerline-only catheter deployment, the mathematically optimal drug release location precisely coincides with the intersection point where the optimal dosing curve begins to deviate from the centerline, providing a definitive, geometry-driven spatial guideline for surgical positioning.

Future research will focus on extending this single-vessel segment framework to complex, multi-branched vascular networks. Recent advancements in machine learning have enabled the highly accurate extraction of whole-brain vasculature geometries [6]. By coupling our stochastic growth model with such precise anatomical network data, we aim to globally optimize targeted catheter treatments within patient-specific whole-organ vascular networks.

## References

[1] S. R. Harrison, et al., Targeted delivery of bioactive molecules for vascular intervention and tissue engineering, Frontiers in pharmacology 9 (2018) 1329.

[2] D. F. Young, Effect of a time-dependent stenosis on flow through a tube, Journal of Engineering for Industry 90 (2) (1968) 248–254.

[3] P. C. Bressloff, J. M. Newby, Stochastic models of intracellular transport, Reviews of Modern Physics 85 (1) (2016) 135.

[4] S. Redner, A Guide to First-Passage Processes, Cambridge University Press, 2001.

[5] O. Bénichou, R. Voituriez, From first-passage times of random walks in confinement to geometry-controlled kinetics, Physics Reports 539 (4) (2014) 225–284.

[6] M. I. Todorov, J. C. Paetzold, O. Schoppe, et al., Machine learning analysis of whole mouse brain vasculature, Nature Methods 17 (4) (2020) 442–449.

